# The dynamics of episodic recall

**DOI:** 10.1101/2025.08.27.671980

**Authors:** Neomi Mizrachi, Ehud Ahissar

## Abstract

The mechanistic understanding of memory recall, a process long known to be influenced by environmental context, has posed a persistent challenge to neuron-centric models. In this study, we approached recall by shifting the focus from internal neuronal representations to brain-environment interactions, employing precise tracking of gaze dynamics within a well-controlled virtual reality (VR) setting. We found that each verbal report of recalled event was consistently preceded by a gradual process of brain-environment coupling: fixational pauses became progressively longer, while gaze direction slowly converged on a specific spatial location associated with the memorized object—a recall-specific location (RSL). Upon the initiation of the verbal report, gaze rapidly diverged from the RSL. Chunked verbal reports were associated with convergence onto shared or spatially clustered RSLs. Moreover, when recall occurred in the same spatial context as encoding, participants recalled objects in the order they were encountered and RSLs were more correlated with gaze locations during encoding. These findings reveal a direct mechanistic dependence of memory recall on the environment and support the view that the environment is not merely a context for memory, but an integral component of the memory itself.

## Introduction

Despite the significant progress in understanding retrieval mechanisms obtained during the last decades, the interaction between recall dynamics and the recall environment remains unclear. It is commonly assumed that retrieving an episode involves the sequential activation of stored neuronal representations of the events, such as object encounters, composing the episode (Atkinson & Shiffrin, 1968; Murdock, 1972; Shiffrin & Raaijmakers, 1992). Existing data support the involvement of dynamic memory traces in the medial temporal lobe and hippocampus, whose activation is followed by the retrieval of individual events (Moscovitch et al., 2016).The retrieval of each such event in the episode is often hypothesized to involve a dynamic convergence to a previously formed attractor (Ackley et al., 1985; Amit, 1989; Hopfield, 1982; Pereira-Obilinovic et al., 2023). In general, such attractors represent low-energy configurations within a dynamic landscape, toward which the system tends to evolve. Indeed, the characterization of memory as stable attractor states has garnered some empirical support (McRae et al., 1997; Ruppin & Yeshurun, 1991), thus far for attractors confined to the neuronal domain.

An embodied view of memory suggests that memory is rooted in motor-sensory processing that emerges from brain–environment interactions (Krakauer et al., 2017; O’Regan & Noë, 2001; Wilson, 2002). In this view, the motor-sensory information encoded during an event becomes part of the memory trace. Consequently, recall involves a recapitulation of motor-sensory interactions (Ianì, 2019; Kent & Lamberts, 2008). Extending the attractor hypothesis to embodied memory implies that a memorized attractor corresponds to a subset of configurations in a multidimensional space spanned by neuronal, motor, environmental, and sensory variables (Ahissar & Assa, 2016). During recall, the system attempts to converge on one of these memorized attractors, guided by the partial information available in the current context (Ahissar & Assa, 2016; Dreyfus, 2002; Merleau-Ponty, 2011). We refer to these embodied attractors as brain–world attractors.

Basing memory on brain–world attractors challenges the notion of memory-specific attractors that exist solely within isolated neural networks—an idea that has also been questioned by substantial empirical evidence showing that such isolation is unlikely. Notably, brain regions associated with memory, such as the hippocampus and entorhinal cortex, have been shown to be functionally connected to oculomotor regions that govern eye movements, particularly the frontal eye field (FEF) and the superior colliculus (SC) (Ryan, Shen, & Liu, 2020).

Oculomotor sensory dynamics include rapid saccadic jumps that bring selected regions of interest into the fovea, and slow drifts that scan those regions during inter-saccadic fixation—or “fixational pause”—periods. Both saccades and fixations have been shown to be closely linked to medial temporal lobe activity (Hoffman et al., 2013; Leonard et al., 2015; Ringo et al., 1994; Sobotka et al., 1997; Sobotka & Ringo, 1997; Staudigl et al., 2017) The propagation of activity between the hippocampus and entorhinal cortex to the oculomotor system is estimated to occur within the duration of a single gaze fixation (Ryan, Shen, Kacollja, et al., 2020). Additionally, hippocampal theta-band oscillations have been shown to be modulated by saccades during visual exploration in ways that predict subsequent recognition (Jutras et al., 2013). Empirical data thus suggest that hippocampal–entorhinal circuits are tightly coupled with oculomotor control. This functional linkage indicates that neuronal dynamics in the hippocampus and entorhinal cortex should not be understood as standalone representations of environmental content. Rather, they may be more accurately viewed as components of broader sensorimotor brain–world attractor systems— dynamical networks that integrate internal neural representations and embodied interactions with the external environment (Linson et al., 2018; O’Regan & Noë, 2001).

Indeed, Hebb (1968), and later Noton and Stark (1971a, 1971b), proposed that gaze behavior is a component of episodic memory. When repeatedly viewing similar drawings, participants tend to produce similar eye movement trajectories (Noton & Stark, 1971a, 1971b). Based on these observations, Noton and Stark formalized the scanpath theory, which predicts that recall is accompanied by the reactivation of eye movements present during encoding. Moreover, the degree of recapitulation of these movements during retrieval is hypothesized to predict recall or recognition performance (Noton & Stark, 1971a, 1971b). Scanpath theory is supported by experimental findings showing that participants tend to fixate on similar locations in the visual field during both encoding and retrieval. This effect has been demonstrated in studies involving the retrieval of scenes (Altmann, 2004), visual objects (Spivey & Geng, 2001), and even auditory information (Scholz et al., 2016), and appears consistent across age groups (Wynn et al., 2018). Furthermore, such recapitulation of eye movements during retrieval improve performance in both recognition (Ryals et al., 2015) and recall tasks (Johansson & Johansson, 2014).

However, the proposed link between memory and the dynamic recapitulation of scanpaths has been questioned (Foulsham & Kingstone, 2013), primarily due to an overreliance on spatial similarity measures alone (Altmann, 2004; Johansson & Johansson, 2014; Olsen et al., 2014; Ryals et al., 2015; Scholz et al., 2016; Spivey & Geng, 2001; Wynn et al., 2018). Moreover, although some studies have examined scanpath similarity in the absence of target objects (Johansson & Johansson, 2014) or with manipulated object configurations (Hannula et al., 2012), the majority have focused almost exclusively on recognition tasks (Brandt & Stark, 1997; Foulsham & Underwood, 2008; Holm & Mäntylä, 2007; Underwood et al., 2009). Recognition tasks typically involve binary judgments—classifying an episode or object as either recognized or unrecognized. Because such judgments can be based on familiarity alone, rather than recollection (Jacoby, 1991; Yonelinas, 1999), recognition tasks may be ill-suited to studying the dynamics of spontaneous, internally cued episodic memory retrieval.

The impact of environmental context on memory recall has been extensively studied, with one robust finding being that memory performance is enhanced when retrieval occurs in a context similar to that in which the information was originally encoded (Godden & Baddeley, 1980; Godden & Baddeley, 1975; Smith, 2001; Tulving, 1972). While several neuronal mechanisms have been proposed for context- or state-dependent memory (Jung et al., 2023; Shulz et al., 2000; Staudigl & Hanslmayr, 2019), the link between context-dependent facilitation and the interaction between eye movements and environmental context remains largely unexplored.

In this study we investigated the temporal dynamics of gaze during the free recall of a rich episodic experience in a virtual reality (VR) setting. Participants performed a free recall task in a naturalistic VR environment, while their head position, eye movements, and gaze trajectories were continuously recorded during both the encoding and recall phases, alongside their verbal reports. The recall environment either replicated the encoding context (“SAME” group) or differed from it (“DIFF” group).

We hypothesized that if object recall involves convergence to a specific brain–world attractor, then participants’ gaze would exhibit convergent dynamics toward object-specific spatial locations (assuming spatial location is one of the encoded features). Moreover, we expected differences in convergence patterns across contextual conditions. In contrast, if memorized objects were not associated with brain–world attractors—or if attractors were limited to internal neural dynamics— we would expect no context- or memory-specific gaze convergence.

## Results

### Convergence dynamics preceding recall report events

Two groups of participants, 30 participants in each, performed an incidental episodic memory test (Sadeh et al., 2018) in the VR. They first had 3 minutes to freely explore and manipulate 15 objects within a virtual room (the “encoding” session, Fig. 1a, left). Then, after a 20-minute break, they were called back into one of two virtual environments and had 3 minutes to recall and verbally report the names of the objects that were present during encoding. The two recall environments were: the same room as during encoding, without the objects (“SAME” context, Fig. 1a, top right) or an environment featuring a white plane and a blue sky (“DIFF” context, Fig. 1a, bottom right). The participants were not informed in advance about the later recall task - they were simply asked to start recalling when the second (“recall”) session started. During both sessions, head, eye and hand movements, along with manual interactions with the objects at the encoding session, and verbal reports during the recall session, were continuously recorded (Fig. 1b-d). In each frame, gaze position was determined by the 3D position of the interaction of the gaze with the environment.

**Figure 1.**
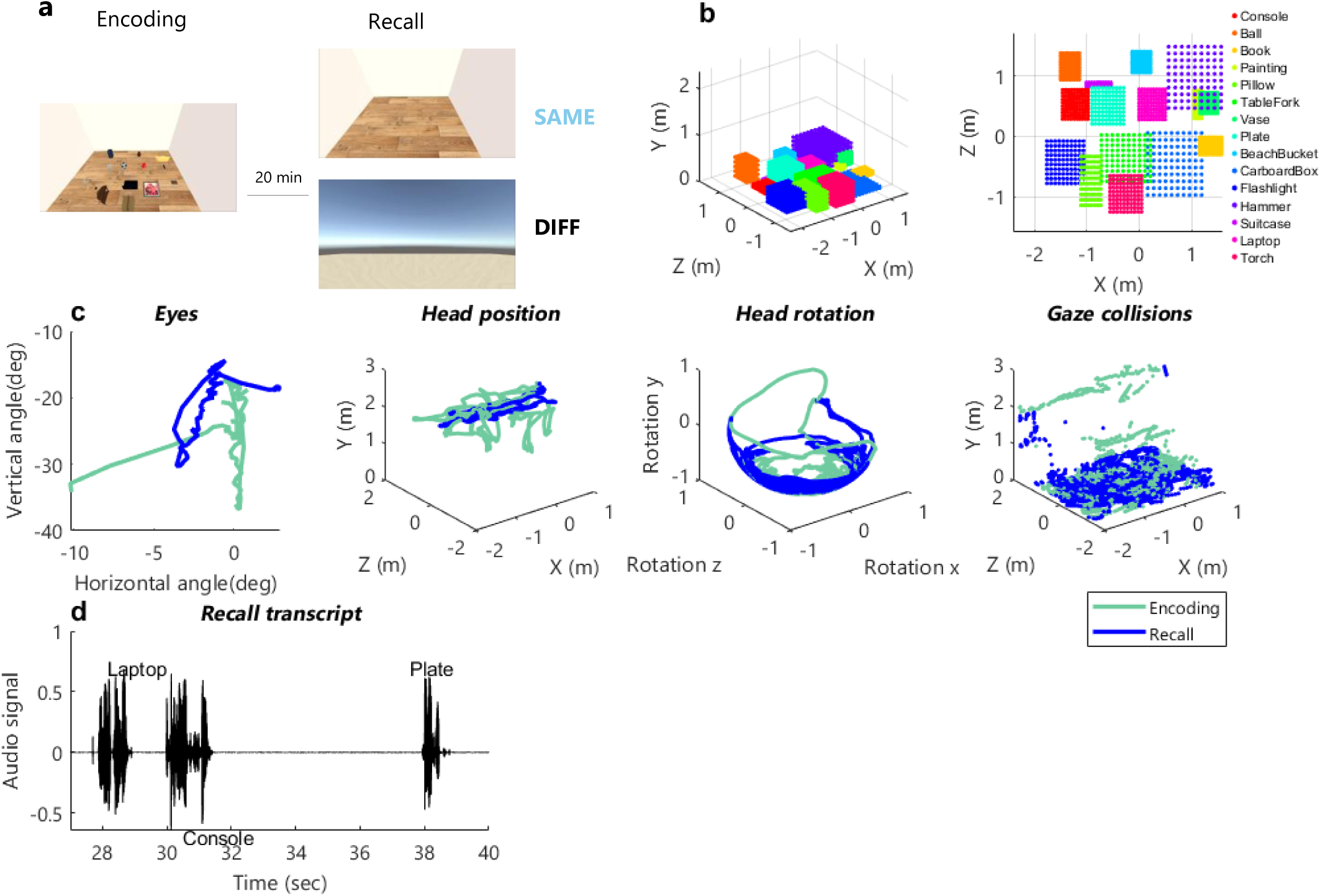
Experimental Design, Virtual Environments, and Data Collection. **a. Experimental procedure and virtual environments.** Left: Encoding environment. Participants freely explored a virtual room containing fifteen everyday objects randomly placed. Right: Recall environments. Twenty minutes after encoding, participants re-entered a virtual room and were unexpectedly asked to recall the names of the objects. Participants were randomly assigned to one of two conditions: the SAME environment (identical to the encoding room but empty), or the DIFF environment (a featureless setting with a blue sky and white plane). **b. Object layout in the encoding environment.** Bounding boxes of the virtual objects in 3D (left) and 2D (right) views indicate their initial locations prior to any manipulation. The dimensions of each box reflect Unity’s *colliders*— defined spatial zones that enable physical interaction with the objects. **c. Tracking data.** Representative example showing eye tracking (a segment of 800 ms is shown), head position, head rotation (in normalized virtual coordinates), and gaze tracking during the entire 3-minute encoding (green) and recall (blue) sessions. **d. Audio data.** Audio waveform of a participant’s verbal recall response.

We first analyzed the dynamics of each participant’s gaze prior to each single recall report event (i.e., the verbal report of a single recalled object name; see Methods). Typical to natural viewing, the participants explored the environment using combined head and eye movements; the trajectories created by the landing positions of the generated saccades are termed “scanpaths” (Noton & Stark, 1971a, 1971b) and the dwelling periods between saccades are termed “fixational pauses” or simply “fixations”.

In both contexts, gaze behavior showed a similar general pattern: consistently, prior to each recall report, the participants’ gaze gradually moved toward a certain location within the virtual environment (Fig. 2). This location, defined by gaze position during the second half of the last fixation before the verbal report, is referred to as Recall Specific Location (RSL) (see Methods;). We analyzed gaze behavior in relation to RSLs and to the time of the verbal report. Our analysis revealed two interleaved dynamical processes.

**Figure 2.**
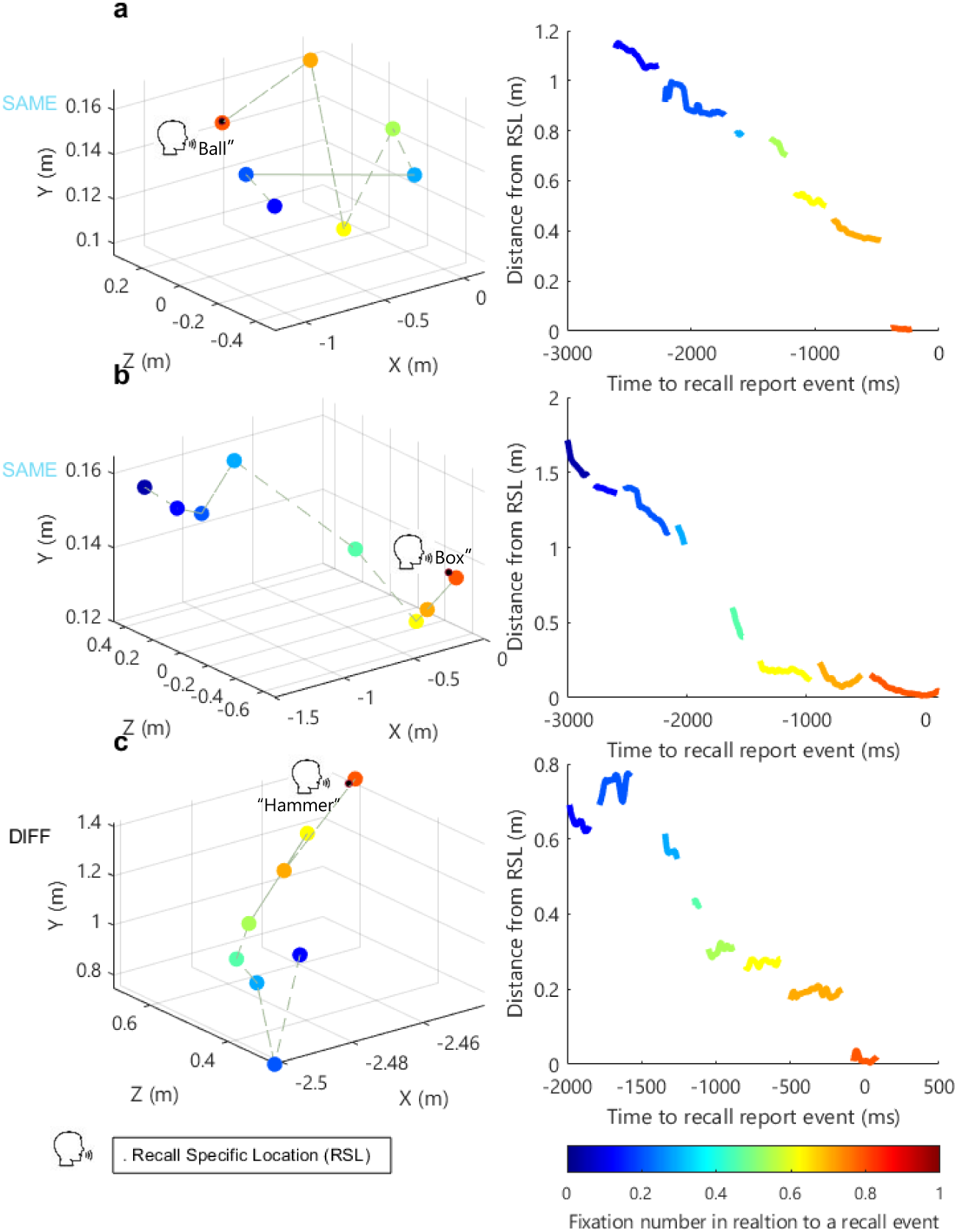
Examples of Gaze Behavior Prior to Recall Reports. Examples of gaze behavior leading up to recall events for multiple participants. **a.** Participant DI from the SAME group, recalling “Ball.” **b.** Participant SSE from the SAME group, recalling “Box.” **c.** Participant OL from the DIFF group, recalling “Hammer.” **All panels:** Left column: Mean fixation positions prior to a specific recall event.. Right column: Distance from each frame in the fixation chain to the second half of the final fixation, defined as the recall-specific location (RSL). Color code indicates the ordinal position of each fixation relative to the recall report.

The first process was a gradual approach to a specific RSL prior to the corresponding verbal report. In both contexts, the distance to the RSL (D) typically decreased gradually during 7 - 10 fixations preceding the recall report (Figs. 2; average number of fixations between events: SAME

= 7.8 ± 3.19 fixations, DIFF = 7.63 ± 3.17 fixations). On average, gaze distance to the RSL gradually decreased in the few seconds leading up to the recall report (Fig. 3a). This trend was also clear when averaging the data for each fixation separately, locked to the onset of the verbal report (Fig. 3b). A similar trend was also observed for gaze angle (Supplementary Fig. 1a). Interestingly, in some cases, movement toward the RSL was observed not only across fixations but also within fixations (Fig. 2 & 3b).

**Figure 3.**
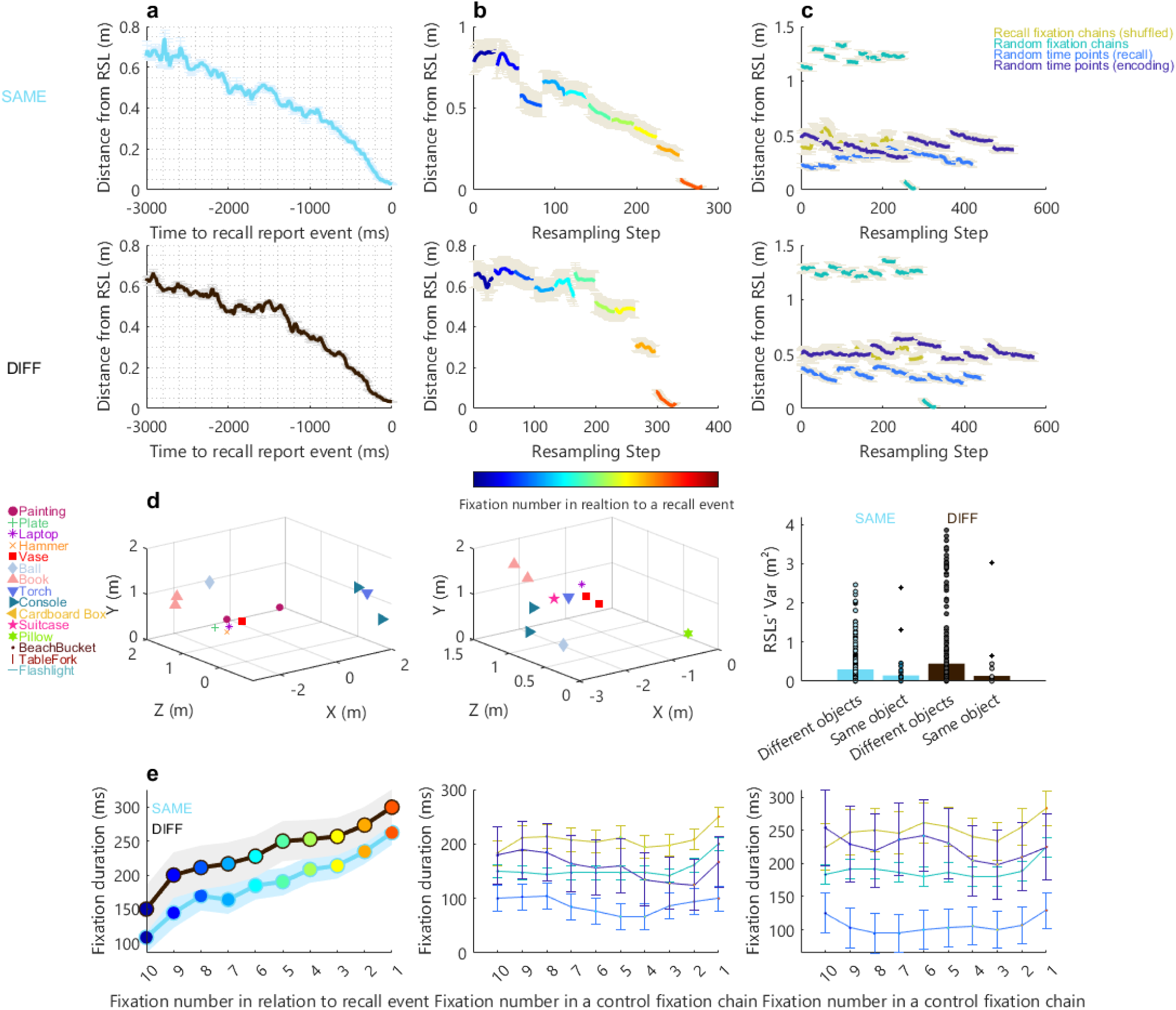
Gaze Behavior Prior to Single-Object Recall Reports. **a.** Distance from the RSL across all single-object recall events for the SAME (top) and DIFF (bottom) groups during the 3 seconds preceding each recall report. **b.** Distance from the RSL across the 10 fixations preceding recall reports. **c.** Distance to the RSL for various control conditions: mixed recall fixation chains (brown), random fixation chains (turquoise), and fixation chains preceding random time points during encoding or recall (light and dark blue). Data are shown separately for the SAME (top) and DIFF (bottom) groups. None of the control conditions exhibited consistent approach behavior (SAME: ρ ≥ –0.5, p ≥ 0.14; DIFF: ρ ≥ –0.5, p ≥ 0.13; Spearman correlation). **d.** Left and middle: RSLs from two participants, each linked to a single-object recall event. Object identity is indicated by colors and shapes. Some objects were recalled multiple times (left: “Book,” “Painting,” and “Console”; middle: “Vase,” “Book,” and “Console”). Note the differing, participant-specific spatial scales. Right: Variability in inter-RSL distances for repeated recalls of the same object versus different objects, shown for the SAME (blue) and DIFF (black) groups. Each data point represents the variance between two RSLs; outliers are marked with diamonds. **e.** Left: Mean fixation duration as a function of fixation number relative to the recall event, for the SAME (blue) and DIFF (black) groups. A strong correlation was observed between fixation duration and fixation number (SAME: ρ = 0.91, p = 2.09 × 10⁻⁴⁰; DIFF: ρ = 0.92, p = 1.57 × 10⁻⁴; Spearman correlation). Middle and right: Fixation durations in the control conditions shown in panel **c** (brown, turquoise, blue, and dark blue), again for the SAME (middle) and DIFF (right) groups. No significant correlations were found (SAME: ρ ≤ 0.23, p ≥ 0.077; DIFF: ρ ≤ 0.2, p ≥ 0.35; Spearman correlation). **Panels b and e:** Colors represent the ordinal position of each fixation relative to the recall report., as indicated in the color bar in panel **b**. **Panels b and c:** Fixations were resampled to a common duration prior to averaging to account for varying fixation lengths (see Methods). **All panels:** Error bars represent the standard error of the mean (SEM).

The relationship between gaze dynamics and the RSL was first evaluated by computing D at each saccade landing position and correlating it with the fixation number (counted backward from the verbal recall report; Fig. 2, Fig. 3b, color-coded from red to blue); D was strongly negatively correlated with fixation number (SAME: ρ = -0.96, p < 5 × 10⁻¹⁵; DIFF: ρ = -0.818, p = 0.006, Spearman correlation). In contrast, no gradual approach toward the RSL was detected in fixation chains preceding random time points or in chains that were randomly generated or randomly ordered (see Methods) (Fig. 3c).

RSLs, approached prior to the recall of different objects, were distributed throughout the recall environment (Fig. 3d). In several cases, participants reported the name of the same object more than once (N = 16 and 9 for the SAME and DIFF conditions, respectively; Fig. 3d, left & middle). Even when these repeated reports were separated by long intervals (SAME: 36.1 ± 11.6 sec; DIFF: 15.02 ± 7.3 sec) they tended to cluster—the variance in RSL positions for repeated reports of the same object was smaller than that for reports of different objects made by the same participant (SAME: 0.14 ± 0.14 m² vs. 0.30 ± 0.40 m²; DIFF: 0.13 ± 0.17 m² vs. 0.44 ± 0.85 m²; bootstrap *p* < 0.005, one-tailed test; Fig. 3d, right). RSLs’ locations within the recall environment and their relation to visual exploration during the encoding session are described in **Context-dependent gaze and report dynamics** below.

The second dynamical process preceding each verbal report was a monotonic, gradual increase in the duration of fixational pauses as the gaze approached the RSL. In the SAME and DIFF contexts, fixation durations gradually increased during the 10 fixational pauses preceding the verbal report (Fig. 3e, left), roughly doubling the duration and yielding a significant correlation between fixation duration and fixation number (ρ >=0.91, p< = 1.57 × 10⁻⁴, Spearman correlation). No such correlation was observed between fixation duration and fixation number in fixation chains preceding random time points, or in chains that were randomly generated or randomly ordered (see Methods) (Fig. 3e, middle and right). On average across all fixations, DIFF fixations were longer (bootstrap, p < 0.005, right-tailed test). In comparison, fixation lengths during the encoding session were not significantly different from one another (SAME: 190.2 ± 8.75 ms, DIFF: 198.01 ± 6.98 ms, bootstrap p = 0.67, two-tailed test).

### Convergence versus divergence from RSLs

In non-linear dynamic systems, convergence and divergence to and from attractors typically follow different trajectories, with divergence being typically faster (Engelken et al., 2023; Pereira & Brunel, 2018). Our data align with this dynamic behavior: in both contexts, the instantaneous speed of gaze toward the RSL (before the recall report event) was significantly slower than the gaze’s speed when leaving the RSL (after the event) (Fig. 4; SAME: toward the RSL: 0.44 ± 1.96 m/s, away from the RSL: 0.62 ± 2 m/s; DIFF: toward the RSL: 0.35 ± 1.79 m/s, away from the RSL: 0.5 ± 1.85 m/s, bootstrap p<0.005, right-tailed test).

**Figure 4.**
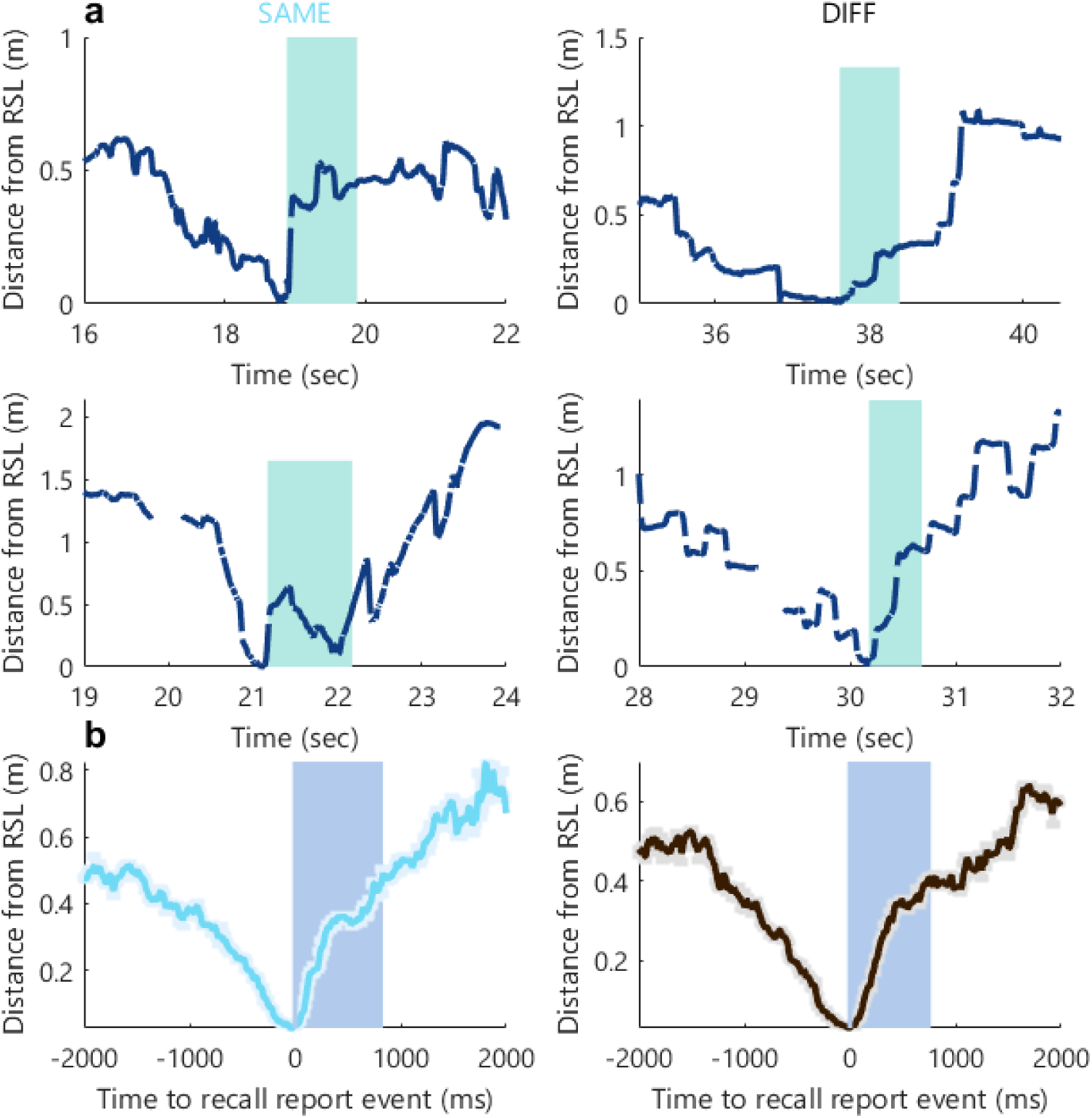
Gaze Dynamics Around Recall Report Onset. **a.** Gaze distance from the recall-specific location (RSL) as a function of time surrounding a single recall event for four participants. gaps indicate blinks. **b.** Median gaze distance from the RSL in the 2 seconds before and after recall onset, shown separately for the SAME (left) and DIFF (right) groups. Error bars indicate the median absolute deviation (MAD). The shaded blue rectangle marks the average duration of verbal recall (SAME: 833 ± 50.93 ms; DIFF: 767 ± 38.79 ms).

Additionally, we found that the rapid shift away from the RSLs occurred typically within a few tens of milliseconds after the beginning of the report (SAME: 30.96±16.5 ms, DIFF:73.47±19.19 ms, bootstrap p=0.07, two-tailed test). This is consistent with the processes of recall and verbal report being distinct (Norman et al., 2019). Specifically, the fact that, in most cases, participants’ eyes did not linger on the location once a verbal report was initiated suggests that the two processes, recall and verbal report, although linked, are not merged. A dynamic consistent with this behavior is that once convergence is achieved at sufficient confidence, a verbal report is initiated and in parallel the gaze system diverges from the reported attractor without waiting for the verbal report to be completed. In the perspective of dynamic systems, the verbal system reports on the dynamic state of the recall process, without being part of the recall closed-loop process.

### Shared or close recall-specific locations (RSLs) and chunking

Recall often fluctuated between single reports (Fig. 5a, purple) and ‘chunks’ of several reports (Fig. 5a, green; see Methods), consistent with previous studies (Romani et al., 2016). We asked whether, in our paradigm, ‘chunking’ depended on the spatial proximity of the relevant RSLs (e.g., Fig. 5b).

**Figure 5.**
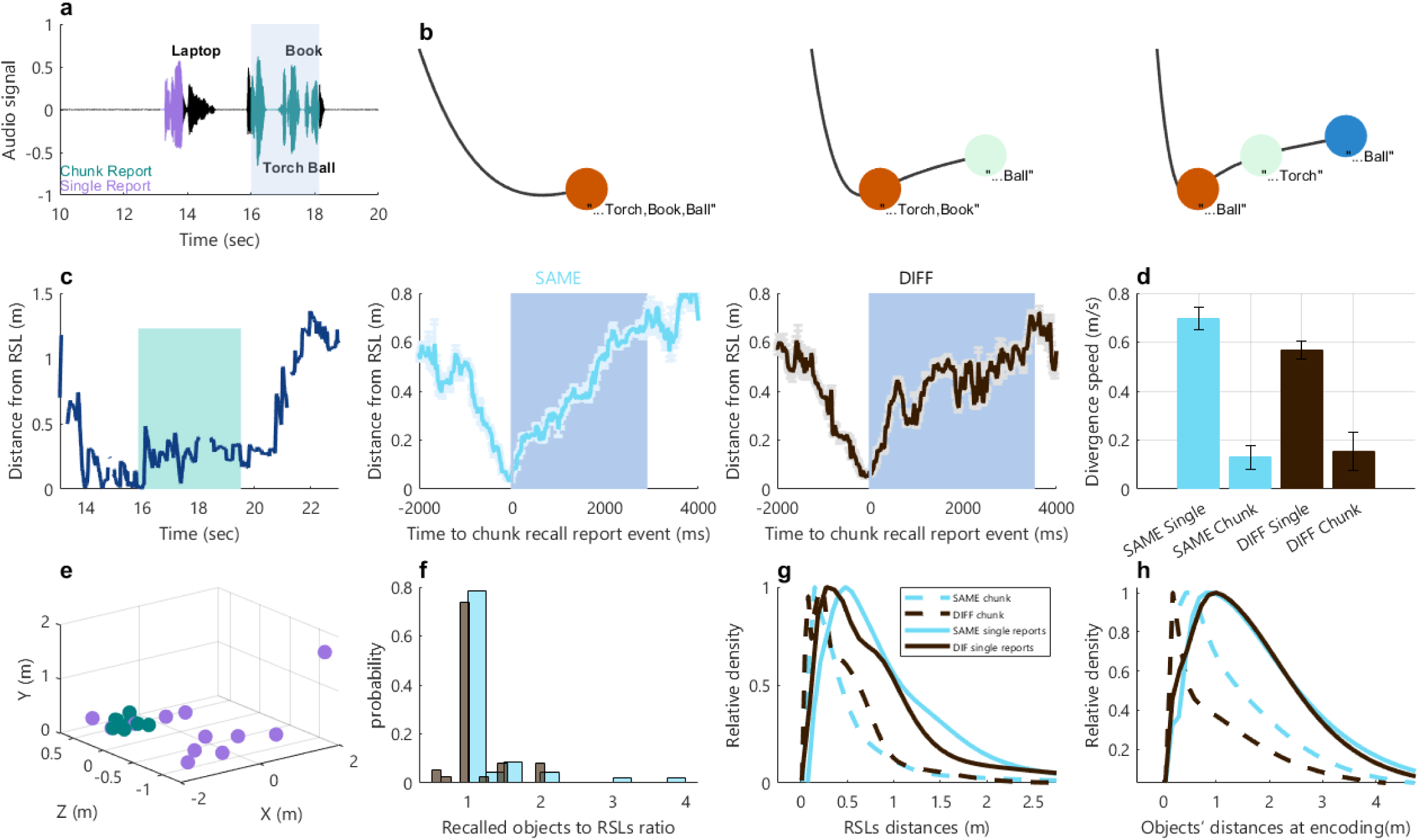
Gaze Dynamics Around Chunked Reports. **a.** A segment of one participant’s recall transcript, with an example chunked report highlighted by a blue patch. **b.** Schematic illustrations of possible chunk–RSL relationships. Left: All objects in the chunk share the same RSL. Middle: Some objects share an RSL, while others have distinct ones. Right: Each object is associated with a unique RSL. **c.** Left: An example of one participant’s gaze distance from the RSL before, during, and after a chunked report (gaps indicate blinks). Middle and right: Median distance to the RSL during the 2 seconds preceding and 4 seconds following chunk report onset, for the SAME (middle) and DIFF (right) groups. The blue patch marks the average duration of chunked reports (SAME: 2904.5 ± 301.18 ms; DIFF: 3537.9 ± 381.1 ms). **d.** Gaze divergence speed (away from the RSL), measured during the verbal report of chunked and single objects in both contexts. **e.** Example from one participant showing RSLs for single-object reports (purple) and chunked reports (green). **f.** The ratio between the number of recalled objects and the number of RSLs within chunk reports for the SAME (blue) and DIFF (black) groups. **g.** Distances between RSLs for objects within a chunk versus consecutive single-object reports, for each group. **h.** Encoding-stage distances between successive chunked objects and successive single objects, shown separately for SAME and DIFF groups. **All panels:** The SAME group is shown in blue; the DIFF group in black. **Error bars:** Median absolute deviation (panel c); standard error of the mean (panel d).

We compared gaze dynamics around chunked report events and single report events. In both contexts, gaze gradually converged on the first chunk object (RSL, Fig. 5c) as observed for single reports (Fig. 4b). Divergence, however, was significantly slower during chunked reports compared with single reports. (Fig. 5c–d; (SAME:0.13 ± 0.047 m/s vs 0.69 ± 0.045 m/s, DIFF: 0.15 ± 0.07 m/s vs 0.58 ± 0.037 m/s; both groups: bootstrap p < 0.005, left-tailed test) (Fig. 5d).

The fact that the gaze did not quickly diverge from the first RSL of the chunk suggests a shared or closely located RSLs between chunked objects. Indeed, in a considerable number of cases, there were more than one report per RSL (Fig. 5b middle and Fig. 5f).

The RSLs included in a chunked report were also spatially closer to each other than those of successive single recall report events (SAME, DIFF: bootstrap p < 0.005, left-tailed test, Fig. 5g). Importantly, this also reflected spatial proximity during encoding; consecutively recalled chunked objects were positioned closer to one another during encoding, compared to consecutively recalled non-chunked objects (Fig. 5h, SAME, bootstrap p =0.0052, DIFF: bootstrap p =0.03, left-tailed test). Interestingly, chunking did not depend on the temporal or sequential proximity between two consecutive object interactions during encoding (SAME: bootstrap p ≥ 0.26, DIFF: bootstrap p ≥ 0.2, two-tailed test).

### Context-dependent gaze and report dynamics

Consistent with previous findings (Smith, 2001), our SAME recall group performed moderately better than the DIFF group. The number of recalled objects did not differ between the groups (Fig. 6a, left). Also, no significant difference was found in the fraction of chunk events out of all recall events (0.29 vs. 0.23, p = 0.18, Mixed-effects logistic regression, F(1, 404) = 1.77). However, participants in the SAME group recalled faster: the duration of their entire recall period was shorter (67.44 ± 41.39 vs. 87.11 ± 40.03 sec, p = 0.02; Fig. 6a, right).

**Figure 6.**
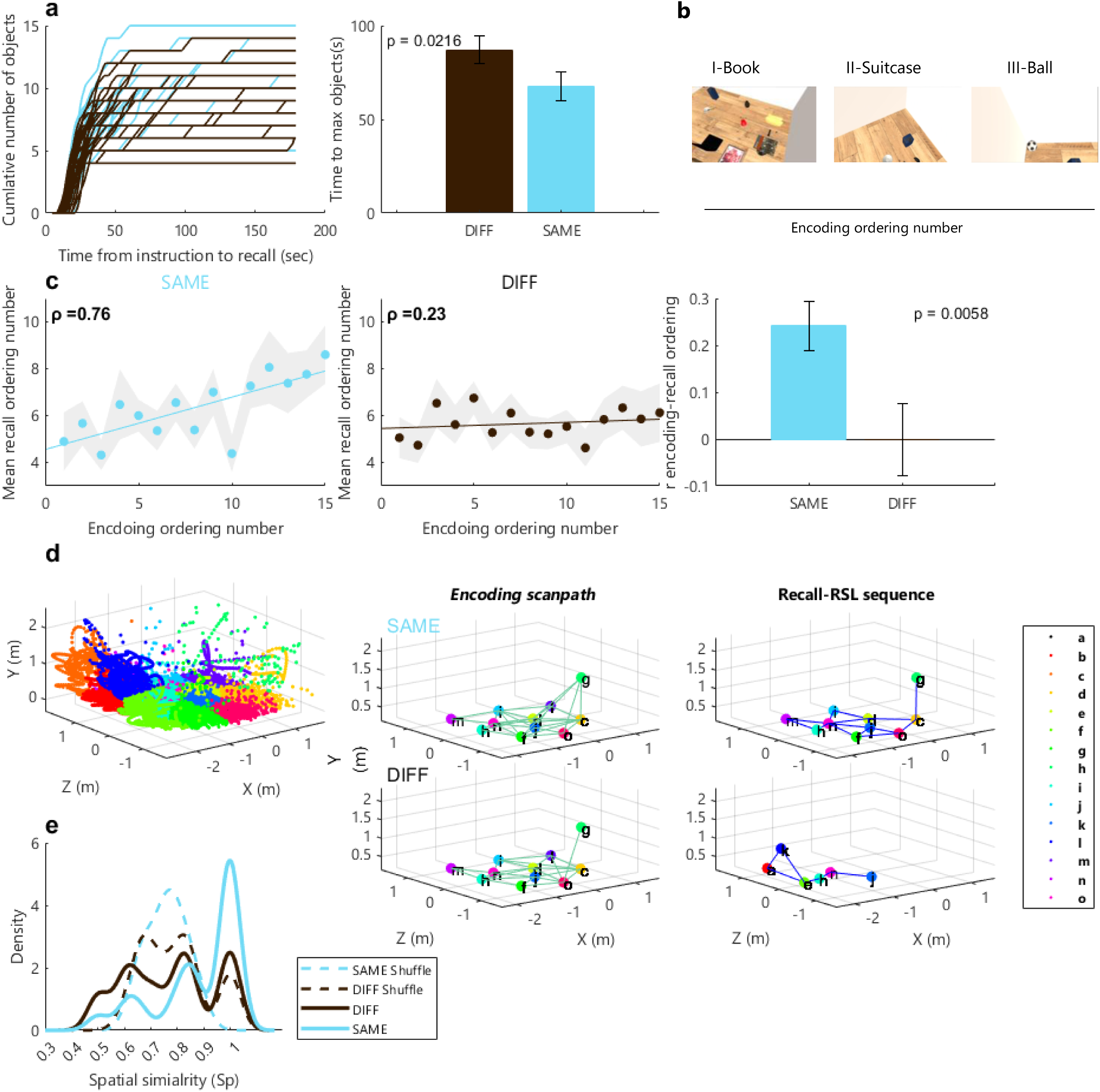
Dependence of Recall on the Spatial Context. **a. Left:** Cumulative number of recalled objects plotted against recall duration. Each trace represents an individual participant from the SAME (blue) or DIFF (black) recall groups. The total number of recalled objects did not differ significantly between groups (SAME: 9.24 ± 2.6; DIFF: 8.93 ± 2.3; *p* = 0.31, one-tailed *t*-test, *t* = 0.47). **Right:** Time to complete object recall for each group. Participants in the SAME group completed their lists significantly faster (SAME: 67.44 ± 41.39 sec; DIFF: 87.11 ± 40.03 sec; *p* = 0.02, one-tailed Wilcoxon rank-sum test, *z* = –2.02, rank-sum = 725). **b.** Example from participant SHM interacting (manually or visually) with three virtual objects (book, suitcase, ball) during encoding. These objects were labeled with an ordinal encoding ordering number (I–III) based on the interaction order. The images show the participant’s view in the virtual environment. **c. Left and middle:** Correlation between average recall order and motor-sensory interaction order during encoding for SAME (left) and DIFF (middle) groups. Each data point represents the mean recall order (averaged across all participants) for objects sorted by their motor-sensory interaction order during encoding. **Right:** Correlation coefficients computed individually for each participant. **d.** Mean fixation positions (across participants and both encoding and recall stages) were grouped into 15 spatial clusters using *k*-means clustering. **Middle and right:** Examples of encoding scanpaths (middle column, green) and recall RSL sequences (right column, blue) from a SAME-group participant (TCH, top row) and a DIFF-group participant (SSE, bottom row). **e.** Probability density estimates of spatial similarity (Sp) indices between encoding scanpaths and RSL sequences, and between encoding sequences and random strings (dashed lines), for the SAME and DIFF groups.

The co-occurrence of shorter fixation durations (Fig.3e, bootstrap, p < 0.005, right-tailed test), and shorter overall recall periods with the SAME context raises an intriguing question. As the overall recall period equals the sum of durations of all individual fixations and saccades during that period, did faster recall with the SAME context result primarily from the fact that fixations were shorter, or were there also differences in the number of fixations? In our data, the average numbers of fixations during recall across all participants were nearly identical in both contexts (SAME: 198.24 ± 5.82, DIFF: 197.85 ± 4.19 fixations, p = 0.78, two-tailed Wilcoxon rank-sum test). Consistently, the duration of the entire recall period was strongly correlated with the sum of the durations of individual fixations (ρ > 0.92, p < 10⁻⁶, Spearman correlation, for both contexts). Thus, faster recall with the SAME context may result from shorter fixation durations.

Unlike with word lists recall tasks, in which object names are provided to the participants, the episode in our experiment was naturalistic, allowing for free movement and unconstrained interactions with objects that were placed in a virtual room. This setup enabled a rich and diverse episodic experience, where objects could be manipulated multiple times, viewed from various angles, or interacted with in combination with other objects (Fig. 6b).

This paradigm revealed an important difference between our two contexts. We quantified the order of motor-sensory interactions with the objects during encoding and correlated it with the order of reports during the recall session. For this analysis we used, for each object, the first motor-sensory interaction with it whether by manually manipulating it or gazing at it during encoding (see Methods). On average, the order of interactions during encoding was strongly correlated with the order of reports during recall in the SAME context (Fig. 6c, left, ρ = 0.76, p = 0.001, Spearman correlation) but not in the DIFF context (Fig. 6c, middle, ρ = 0.23, p = 0.4, Spearman correlation). The same contrast between contexts was observed when these correlations were computed for individual participants (Fig. 6c, right, SAME: 0.24 ± 0.05, DIFF: - 0.009 ± 0.07, Spearman’s ρ, p = 0.005, right-tailed t-test, t = 0.08).

We next compared the similarity of encoding and recall gaze dynamics by creating fixations’ clusters (Fig. 6d), as used in previous studies (Brandt & Stark, 1997; Duchowski et al., 2010), using an approach similar to that used by Wynn (2016). A data-driven definition of the region of interest was used to create 15 clusters using the mean position of all participants’ fixations (see ‘Methods’, Fig. 6d, left). The mean position of fixations (in encoding or recall trajectories) was replaced with the cluster notation (Fig. 6d, middle, right). Using these clusters, we created the encoding scanpath (comprising all encoding fixations) and the RSL sequence (comprising the fixations that were used to define the RSLs). We then evaluated the spatial similarity between the encoding scanpath and recall RSL sequence by a spatial similarity index (Sp), calculated as the number of shared clusters divided by the total number of clusters in the RSL sequence.

The number of visited clusters was similar across groups during encoding (SAME: 11.3 ± 1.0; DIFF: 11.8 ± 1.3) and recall (SAME: 6.6 ± 2.1; DIFF: 6.3 ± 1.6) (*p* > 0.3; Wilcoxon rank-sum test). However, Sp scores were significantly higher in the SAME context (Fig. 6e, SAME: 1 ± 0.12, DIFF: 0.80 ± 0.14; values are median ± MAD; *p* = 0.009, rank-sum = 677.5, *z* = 2.33; right-tailed Wilcoxon rank-sum test). Notably, only in the SAME group the number of recalled objects correlated with the fraction of clusters from the encoding scanpath that reappeared in the RSL sequence (SAME: ρ = 0.49, *p* = 0.01; DIFF: ρ = –0.09, *p* = 0.61; Spearman correlation; adjusted α = 0.025 [0.05/2]). Additionally, only in the SAME context, Sp scores were significantly higher than those of randomly generated strings (Fig. 6e, SAME: *p* = 0.002, signed-rank = 137, *z* = 2.80; DIFF: *p* = 0.70, signed-rank = 44; right-tailed Wilcoxon signed-rank test).

## Discussion

We used a free recall task in virtual reality environments to explore the dynamics of memory retrieval in context-dependent episodic memory. We asked whether recall arises through convergence to brain-environment steady states. We found consistent patterns: slow gaze convergences toward specific object-related locations (RSLs) prior to recall verbal reports, rapid divergences after triggering a verbal report and chunking of objects in nearby RSLs. When recalled in the same context as during encoding, objects were retrieved in an order correlated with the order of their interactive encoding. We propose that these patterns are consistent with the view of episodic memories as brain-world steady states, or attractors.

Identifying an attractor in a dynamical system is a challenging task. One criterion that should be satisfied is that the system’s dynamics should be found to prefer a subset of states that corresponds to the attractors (Khona & Fiete, 2022). The RSLs observed here may represent such a subset: prior to recall verbal reports, gaze progressively converged to the RSLs, which form a subset of all possible positions. Simultaneously, fixation durations steadily increased, with the final fixation before a recall report being the longest (Figs. 2 & 3), further emphasizing the preference of the system to dwell nearby RSLs.

Attractor states are also expected to persist over time and after removal of their relevant input and be invariant across conditions and across behavioral states (Khona & Fiete, 2022)). Our results align with these principles: attraction to RSLs occurred long after their relevant sensory input was removed, and under conditions that differed from those existing during encoding. A further test of attractor invariance is whether different starting points lead to the same RSL when reporting the same object. Indeed, in the subset of trials in which participants reported the same object name more than once, the variance in RSLs of the repeated reports was significantly smaller than the variance between RSLs of different objects (Fig. 3d), demonstrating spatial consistency.

Another key feature of attractor dynamics is resilience to perturbation: the system should return to the attractor following a perturbation (Khona & Fiete, 2022). Our data partially support this prediction as well. In chunked reports, after leaving an RSL, the gaze system was often drawn back to the same or a spatially nearby RSL. If individual object reports within a chunk are treated as perturbations, this pattern may reflect the system’s being re-attracted to the chunk’s attractor. Overall, thus, our results are consistent with the description of episodic memories being encoded in steady-states, or attractors, of the dynamic system that includes both the brain and its environment. Additional perturbation tests can be applied in future experiments to further explore this point. Such tests can include complete darkening of the environment for varying durations or presentation of brief distractors during convergence or while gazing at an RSL.

Our participants were asked to retrieve object names without explicit instructions to recall their locations. Nevertheless, they consistently used spatial information through gaze dynamics, linking RSLs to object names. This aligns with previous research suggesting that spatial information is automatically encoded and used to organize recall (Mandler et al., 1977; Robin et al., 2016), that head orientation-related activity is reactivated during successful retrieval (Schreiner et al., 2024), that participants reenact eye movements even when asked to recall auditory information (Scholz et al., 2016), and that miniature eye movements can reveal visual-spatial information (Linde-Domingo & Spitzer, 2024). Moreover, theta oscillations, which have been shown to encode spatial information and segment navigational routes, also support imagined navigation (Seeber et al., 2025). In our experiment, the connection between specific memories and their associated spatial location is further evidenced by the chunking of object reports: objects recalled together frequently corresponded to the same or adjacent RSLs (Fig. 5).

Spatial location is likely one of many dimensions that make up memory-related attractors. Other relevant dimensions spanning the space in which episodic memories are retrieved are dimensions based on neuronal activity in the visual system, the hippocampus and hippocampus-associated medial temporal lobe areas, such as the entorhinal and perirhinal cortices. If convergence to RSLs involve both gaze and MTL dynamics, their dynamics should exhibit similar time scales. In our experiments gaze spatial convergence occurred within 7 - 10 fixations, or ∼2 - 3 seconds. This time scale indeed aligns with the time scales described for neuronal processes accompanying recall events. In free recall experiments employing response-locked analysis, hippocampal activity was found to precede recall reports by ∼1 - 2 seconds (Gelbard-Sagiv et al., 2008; Norman et al., 2019). Recall-related interactions between hippocampal and cortical activities last between ∼2 to 3 seconds (Staresina & Wimber, 2019).

Additional evidence for a link between oculomotor behavior and hippocampal dynamics comes from findings that sharp wave ripple events co-occur with visual, goal-directed exploration (Leonard et al., 2015; Leonard & Hoffman, 2017). These sharp wave ripple events were associated with longer fixations (Leonard et al., 2015). Sharp wave ripples rate was also shown to increase before a recall report (Norman et al., 2019). Taken together with our results (Fig. 3e), these observations suggest that sharp wave ripples may be a part of an MTL-gaze closed-loop convergence process toward memorized brain-world attractors.

Beyond their temporal alignment with MTL patterns, the observed gaze dynamics also satisfy a key prediction of attractor dynamics: convergence into an attractor is typically slower than divergence from it (Engelken et al., 2023; Pereira & Brunel, 2018). The dynamics around RSLs observed here satisfies this prediction as well - typically, convergence to an RSL was significantly slower than divergence from it (Figs. 4b). Notably, divergence often began in parallel with the onset of the verbal recall, suggesting that reaching the attractor triggers the recall response.

We further investigated the idea of memories as brain-world steady-states by testing recall in the different environmental contexts. During recall, some of the environmental aspects are missing. If indeed memories are stored as attractors of these dynamic systems, then it means that recall dynamics depend on the amount and quality of the available environmental aspects. Consistent with this, we found that both gaze behavior and recall dynamics were influenced by contextual information.

Context-dependent memory has long intrigued researchers, with studies showing that people often perform better in recall and recognition tasks when the context matches that of the original learning environment (Godden & Baddeley, 1980; Godden and Baddely, 1975; Smith, 2001; Tulving, 1972). Our findings are in line with these studies, with the SAME group exhibiting faster retrieval of object names. In addition, our data reveal that the availability of sensory information also impacts the temporal order of retrieval (Fig. 6c). It has been proposed that memory recall unfolds in the same order as the original experience (Howard & Kahana, 2002; Kahana, 1996; Michelmann et al., 2019). Our results indicate that this process depends strongly on the availability of contextual sensory information.

While previous studies have demonstrated that reenactment of encoding eye movements occurs during subsequent views of the same stimuli (Brandt & Stark, 1997; Foulsham & Underwood, 2008; Holm & Mäntylä, 2007; Underwood et al., 2009) or even in the absence of the stimuli, as observed in the ‘looking at nothing’ phenomenon (Martarelli & Mast, 2013), a direct comparison of eye movement reenactment across different contextual conditions has hitherto remained unexplored. Our results demonstrate that gaze behavior differ significantly between the two contextual conditions.

In both conditions, gaze converged on object-specific RSLs that were spatially related to visual exploration during encoding (Fig. 6). However, under SAME recall conditions, these RSLs were more closely aligned with the original exploration locations, and the proportion of encoding scanpath clusters that reappeared in the RSL sequence predicted participants’ recall performance. The presence of encoding-related spatial context also allowed faster visual processing during individual fixations (Fig. 3e). The tight relation between mean fixation duration and overall duration of the recall period, together with the fact that the number of fixations during the recall period was similar across conditions, suggests that the fixation may serve as a meaningful processing unit during episodic recall: recall may terminate once a sufficient number of fixations have been employed, with context-specific processing unfolding within each fixation (Rucci et al., 2025). This hypothetical idea warrants careful empirical investigation.

Our findings are thus consistent with the idea that memorizable brain-world interactions are formed during each episode. Our data further suggest that during recall, brain-world interactions are attracted toward those memorized attractors. Successful convergence on an attractor triggers a verbal report of its label. This research extends the concept of attractor-based memories, suggesting that they are not solely based on neuronal representations but are grounded in brain– world dynamics.

## Methods

### Experiment

#### Participants

The SAME group consisted of 30 participants (14 females, aged 19-37, M = 27.4 ± 4.9 years), and the DIFF group included 30 participants (14 females, aged 18-32, M = 25.01 ± 3.74 years). All participants were naïve to the task and provided written consent in accordance with the approved Weizmann Institute Review Board (IRB) guidelines. All participants were healthy, with no visual or motor impairments, and were native Hebrew speakers or completely fluent. The effective sample size was constrained by the quality of the eye-tracking data (see "Eye Tracking Data Processing") and the virtual scene display, with variation between participants. The sample sizes for each session were as follows: Recall session gaze analysis (SAME = 25, DIFF = 28), Recall performance analysis (SAME = 29, DIFF = 29), Encoding session analysis (SAME = 26, DIFF = 29), and the gaze similarity between the Encoding and Recall sessions (SAME = 22 DIFF = 28).

#### Sample size and power estimation

The required sample size for this experiment was estimated based on the results of a pilot study, which included 10 participants per experimental group. Initial analysis of the pilot data allowed us to estimate the necessary sample size based on the observed effect sizes. Sample size calculations were performed using G*Power software, with an alpha level of 0.05 and a power level of 0.95. The effect sizes indicated that 12 to 29 participants would be required per group. We chose to include 30 participants in each group to ensure sufficient power for the analysis.

#### Experiment design

The experiment consisted of two sessions - ‘encoding’ and ‘recall’ - and included two participant groups. During the encoding session, participants from both groups were asked to freely explore a Virtual Reality (VR) room with virtual objects (Fig. 1a, left). Twenty minutes after encoding (following the procedure outlined in (Sadeh et al., 2018), participants entered one of two recall environments: ‘SAME’ (the same virtual room, but empty of objects, Fig. 1a, top right) or ‘DIFF’ (an environment that did not resemble the encoding environment, Fig. 1b, bottom right). In the recall session, participants were unexpectedly asked to recall the names of the objects from the encoding environment.

#### Experiment procedure

After providing informed consent, participants were introduced to the virtual reality (VR) Head-Mounted Display (HMD). Following this, the eye tracker was calibrated to each participant’s specifications. Participants were then briefed on how to use the controllers to interact with virtual objects, with demonstrations on how to grab (using the grip button) and drop (using the trigger button) objects.

Next, participants entered the *Practice environment* (see ‘Virtual Environments’), where they were asked to practice grabbing and dropping objects. After one minute, they were asked whether they understood how to use the controllers. If they responded negatively, they repeated this practice session until they were comfortable with the controllers.

Once familiar with the controllers, participants were informed that they would soon enter the *Encoding Environment* (Fig. 1a, left, see ‘Virtual Environments’), where they could freely explore a virtual room with objects. The encoding session lasted for 3 minutes. A 15-minute break followed the encoding session, after which eye-tracker calibration was repeated.

Twenty minutes after the encoding session, participants entered one of the two recall environments: ‘SAME’ or ‘DIFF’ (Fig.1a, right, see ‘Virtual Environments’). Before entering the recall environment, participants were informed that they would hear task instructions through the HMD headphones and were asked to perform the task to the best of their ability. The recall session lasted for 3 minutes.

#### Virtual environments

Four virtual environments were created using Unity 2019.4.

##### Practice Environment

Three virtual objects—a cylinder, a capsule, and a rod—were randomly placed in a virtual room measuring 4 x 2.2 x 3.3 meters. These objects could be grabbed and dropped, similar to the ‘*Encoding environment’* objects*. Encoding Environment*: A virtual room, also sized at 4 x 2.2 x 3.3 meters, was created. Fifteen everyday virtual objects were randomly placed in the room (Fig. 1a, top, left): ‘Console’, ‘Ball’, ‘Book’, ‘Painting’, ‘Pillow’, ‘Table Fork’, ‘Vase’, ‘Plate’, ‘Beach Bucket’, ‘Cardboard Box’, ‘Flashlight’, ‘Hammer’, ‘Suitcase’, ‘Laptop’, and ‘Torch’. The objects were randomly selected from a set of virtual items from the SteamVR asset library. None of the objects shared a Hebrew name with another. All objects were everyday items to facilitate common interactions and actions. Participants could grab and drop these objects. Grabbing was enabled by pressing the controller’s grip button when participants were close enough to the object’s center (≤ 0.35 m). Dropping was achieved by pressing the trigger button. Objects could be held simultaneously with both the left and right controllers, and multiple objects could be held with each controller*. SAME Recall Environment:* Was identical to the encoding environment, except it was empty of objects*. DIFF Recall Environment:* featured a white plane and a blue sky. In both recall environments, an instruction to recall the names of the objects was played through the HMD headphones, 5 seconds after entering the environment. Participants’ audio reports were recorded using the HMD embedded microphone.

#### Head-mounted display (HMD)

A VR-based head-mounted display (HTC VIVE Pro Eye, HTC Corporation) was used in the experiment. The HMD features a refresh rate of 90 Hz, a resolution of 1440 x 1600 pixels per eye, and a field of view of 110 degrees.

#### Tracking of eyes, head, and hands

Head and hand movements were tracked at 90 Hz using the StemVR tracking sensors embedded in the HMD and controllers, along with the SteamVR tracking system. Eye movements were tracked using the video-based SR Anipal eye tracker integrated into the head mounted display. The eye tracker was used in conjunction with its SDK (version 1.3.1) and SR Runtime (version 1.3.2). As previously reported (Imaoka et al., 2020), consistent 120 Hz tracking data could not be achieved with the eye tracker, so we resampled the data (see "Eye Tracking Data Processing"). The following parameters were recorded from both eyes: gaze origin (in mm, relative to the head center, along the x, y, and z axes), gaze direction (normalized between 0 and 1, along the x, y, and z axes), pupil diameter (in mm), eye openness, and data validity (in normalized units). Eye tracker accuracy was between 0.5° and 1.1° (Imaoka et al., 2020). Calibration and validation were performed using five points, and the eye tracker’s software was used for these procedures.

### Data analysis

Eye Tracking Data Processing: Eye tracking data were collected throughout all sessions of the experiment (see ‘Tracking of Eyes, Head, and Hands’). As previously reported (Imaoka et al., 2020), we were unable to sample the eyes at the intended 120 Hz rate; instead, the sampling intervals varied, with a mean between-frame interval of 11.46 ± 40.7 ms, compared to the expected 8.33 ms. The eye tracking data were then resampled to a consistent frame rate of 120 Hz using the MATLAB function ‘*resample*’ with the *‘spline’* option. Only frames with valid eye measurements (validity = 31, indicating accurate tracking of eye openness, pupil position, and direction across all axes) were retained. This definition automatically excluded blink frames (Imaoka et al., 2020). Additionally, frames with unrealistic pupil values (<1.5 or >5.5 mm) and those preceding or following such frames (identified by points of maximum acceleration within a 30-frame window, or 250 ms) were removed. Frames with missing tracking (time jumps exceeding one second) were also discarded. Only participants with at least 80% valid frames in a specific session were included in the analysis of eye movements and gaze. The fraction of valid frames for participants included in the eye movement analysis was as follows: SAME: encoding = 0.93 ± 0.12, recall = 0.87 ± 0.12; DIFF: encoding = 0.92 ± 0.12, recall = 0.88 ± 0.05. Only the dominant eye was analyzed.

#### Categorization of eye movement trajectories into saccades and drifts

Eye movement trajectories were categorized into saccades and drifts using a velocity-based algorithm (Bonneh et al., 2010). The data were first smoothed using a Savitzky-Golay filter with an order of 3 and a window size of 11 (MATLAB ‘*sgolayfilt*’). The following parameters were used to differentiate saccades from drifts: segments where the minimum velocity exceeded 28 deg/sec, peak velocity exceeded 40 deg/sec, the amplitude was at least 1 degree, and there were no angle changes greater than 15 deg/sec.

#### Combination of head and eye trajectories into gaze

During the experiment, head and eye trajectories were continuously combined to create a gaze ray, with a specific origin (calculated using Unity’s *‘TransformOrigin’* function) and direction (calculated using Unity’s *‘TransformDirection’* function). The collisions (position in meters) of this gaze ray with virtual objects and environment colliders in the Unity scene (see ‘Virtual Environments’) were calculated using Unity’s ‘*RaycastAll*’ function, and the corresponding timestamps were recorded.

Since head and eye movements were recorded at different sampling rates, we resampled the head trajectory to 120 Hz post-experiment and aligned the head and eye data offline for gaze calculation. We then recalculated all gaze collisions using these aligned trajectories, which were subsequently used in the analysis.

Since gaze-ray collisions depend on the colliders in the scene and the objects within it, and because scene colliders varied across different virtual scenes (with different configurations; Fig. 1a), as well as the movable positions of virtual objects during the encoding session (Fig. 1a, left), we aimed to standardize collision calculations across all scenes. Specifically, we sought to ensure that a gaze ray with the same origin and direction would yield consistent collision points (measured in meters) regardless of scene or object configuration. To achieve this, we computed the trajectory of gaze collisions within the SAME recall environment, and these collision points were used in all analyses presented here, except for the output position curve, where collisions on the actual object’s surface were utilized (see ‘Creation of Output Position Curves’).

#### Transcription of audio files

Each recall recording was transcribed using Google’s Speech-to-Text API. The resulting transcription included the pronounced Hebrew words along with their corresponding start and end timestamps (Fig. 1d). Each word was assigned to a specific object name or categorized as ‘Description’ (descriptive adjectives for objects, e.g., “small,” “red”), ‘False Memory’ (referring to non-existent objects), or ‘Other.’ If a time interval contained multiple words (referred to as ‘chunk report events’), the interval was equally divided among the words, with each word assigned its own start and end timestamps. Transcriptions that included only a single object name are referred to as "single recall reports," while those containing multiple words within a chunk event are referred to as "within-chunk reports." Some transcriptions included a rapid repetition of the same object name twice (SAME: 13, DIFF: 8 events), initially recognized as chunks. In these cases, the first pronunciation of the object was included in the single recall report analysis, while the second pronunciation was discarded.

#### Association of fixation chains to specific recall reports

Gaze collisions (see Methods, ‘Combination of Head and Eye Trajectories into Gaze’) of fixations preceding each recall report event were analyzed. Only fixations that both preceded the current recall report event and started after the previous one were considered, with a maximum of 10 fixations included. Only recall events in which the final fixation lasted at least 50 ms were included. This resulted in the following number of fixations’ chains: single report events: SAME: 124, DIFF: 135; chunk report events: SAME: 46, DIFF: 38.

#### Recall Specific Location (RSL) definition

A Recall Specific Location (RSL) was defined for each fixation chain that preceded a recall report event. The RSL was calculated as the mean position of the last half of the final fixation before a recall report. We used this calculation rather than relying on the objects’ absolute positions, as participants often interacted with the objects multiple times and in different locations during encoding.

#### Distance or angle to the RSL

The distance or angle to the RSL was computed for gaze trajectories (including both saccades and fixations) for each frame, 2 or 3 seconds before recall report initiation (for both single report events and chunk report events) and 2 or 4 seconds afterward (for the single and chunk reports, correspondingly).In addition distance or angle were computed using fixations only in chains preceding recall reports. To account for fixations of varying lengths, we resampled all fixations to their average length before averaging across fixations (length resampled to SAME = 28, DIFF = 33 sampling steps). The distance or angle was smoothed using a moving average with a window size of 3 frames.

#### Comparison of fixation chains with control conditions

We compared the slopes of fixation chains prior to recall events to those of four control conditions: (i) Fixation chains prior to recall events that were mixed into a random order (SAME, n=111; DIFF, n=130 chains). Not all recall events were used (SAME=124, DIFF=135) because some first fixations in the remixed chains did not meet the length requirement (≥50 ms). (ii) Random fixation chains of 10 fixations, pooled from the recall trial (SAME, n=850; DIFF, n=900 chains). (iii) Fixation chains selected prior to random time points during the encoding or recall trials (SAME, n encoding=1909; n recall=1908; DIFF, n encoding=2024; n recall=2111 chains). Control conditions fixation chains were resampled to their average length to allow averaging across fixations (SAME: 28-52, DIFF: 33-56). The correlation between the number of fixations in relation to a recall report event and the distance to the RSL were calculated for both real and control events.

#### Comparison of RSLs of repeated recall vs. RSLs of different object recall

The distance between RSLs were quantified as the mean variance of their positions across the x, y, and z axes. The mean variance was compared between the RSLs of repeated reports (N = 16 for the SAME group and N = 9 for the DIFF group) and the variance between the RSLs of different objects recalled by the same participant. For each participant, the mean RSL variance of different objects was estimated between all possible object pairs (SAME=342 pairs, DIFF=351 pairs). Outliers were detected and removed using the MATLAB function ‘rmoutliers’.

#### Gaze velocity toward or away from the RSL

Instantaneous gaze velocity toward or away from the RSL was estimated by smoothing the gaze distance with a moving average (window size = 5) and computing the absolute value of its derivative.

#### Creation of output position curves

To compute the output position curves, it was necessary to assign an encoding ordering number to each object. During the encoding session, each frame in the trial was labeled with the name of the object that was either being looked at or manipulated. Whether an object was visually attended to was determined by gaze collisions with the object (see "Combination of Head and Eye Trajectories into Gaze"). Object manipulation was identified based on the time between object grabbing and dropping, as recorded by the Unity software. Each frame received a label corresponding to the object being interacted with. If an object was interacted both visually and manually, or with only one sense (gaze or hand manipulation), the frame was labeled with that object’s name. Frames where different objects were interacted with by hand and gaze simultaneously were discarded. For objects that were interacted with multiple times, only the first interaction was used to determine the encoding order. The encoding order number was then correlated with the order in which object names were reported during the recall session.

#### Creation of the encoding scanpaths and recall RSL sequences

To create the encoding scanpath and recall RSL sequence, we pooled the mean locations of gaze fixations lasting at least 50 ms across participants and groups. We collected all encoding fixations and the last fixation before the recall report events (the same fixation used to define the Recall Specific Location (RSL), for both single reports and chunk reports). To enable recall fixations to contribute to cluster formation similarly to encoding fixations, despite their markedly different frequencies (all encoding fixations were included, whereas only fixations preceding recall reports were considered for evaluating RSL sequences during recall), and to ensure equal contribution of all subjects to cluster formation, we resampled the fixation trajectories to a consistent length of 215 points. After resampling, we pooled all equal-length trajectories and clustered each point (n=21070 points) into one of 15 groups using k-means clustering (Fig. 6d, left). The initial cluster locations were based on objects’ initial positions during encoding. This data-driven definition of regions of interest (Wynn et al., 2016) is believed to better mitigate resolution and saliency effects (Wynn et al., 2016). Each fixation mean position was then assigned to one of the clusters, and the encoding scanpaths and recall RSL sequences were subsequently generated (Fig. 6d, middle, right).

#### Similarity of encoding scanpaths and RSL sequences

To assess the similarity between encoding scanpaths and recall RSL sequences, following an approach used in previous studies (Duchowski et al., 2010; Privitera & Stark, 2000) we computed the following similarity index:

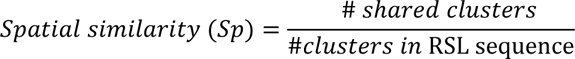

The similarity between each subject’s encoding scanpath and their recall RSL sequence was compared to the similarity between the encoding scanpath and 1,000 randomly generated strings of the same length (215), each containing the same number of unique clusters as the subject’s RSL sequence.

#### Statistical analysis

Each pair of compared distributions was tested for normality using the Anderson–Darling test. For comparisons of independent distributions, if at least one distribution was identified as non-normal, the Mann–Whitney test was used; for paired comparisons, the Wilcoxon signed-rank test was applied. Otherwise, if both distributions were normal, a t-test (for independent distributions) or a paired t-test (for paired comparisons) was used. Correlation between ranks was assessed using Spearman’s correlation.

For comparisons involving dependent samples (e.g., the derivative of gaze before and after recall reports, proximity of RSLs or object positions during chunk or single report events), bootstrap methods were applied. In this procedure, samples from both groups were pooled, and 10,000 iterations were performed. In each iteration, two samples were drawn from the mixed pool—one sample the size of group I and the other the size of group II. The difference in the median values of the two samples was calculated for each iteration. The fraction of bootstrap values more extreme (either larger or smaller, depending on the case) than the experimental value was used to compute the probability of obtaining the experimental result by chance.

The proportion of the binary dependent variable was analyzed using mixed-effects logistic regression with subject-level random intercepts. Coefficients representing the model’s fixed effects were tested for significance using ANOVA and the F test.

The use of one-sided or two-sided tests was chosen according to tested hypotheses and is mentioned in the text. Corrections for multiple comparisons used the Bonferroni method.

#### The use of large language models (LLMs) for text editing

In this study, ChatGPT-4 was employed in few cases to verify syntax and grammar.

## Supporting information

Supplementary Figure 1. Supplementary material accompanying Figure 3.

## Experiment setup

The entire virtual reality setup used to run this experiment is freely available at: https://zenodo.org/records/16901726

## Data availability

The data used in this study are freely available at: https://zenodo.org/records/16901726

## Code availability

The custom code used in this study is freely available at: https://zenodo.org/records/16901726

## Acknowledgments

We thank Yoni Pertzov, Alexander Rivkind, Michele Rucci and Itzik Norman for their invaluable feedback and insightful comments on earlier drafts of the manuscript. This project has received funding from the European Research Council (ERC) under the EU Horizon 2020 Research and Innovation Programme (grant No. 786949), the USA Air Force Office of Scientific Research (AFOSR, grant No. FA9550-22-1-0346) and a research grant from the Estate of Thomas Gruen.

