## Supplementary Figure 1. Supplementary material accompanying Figure 3. for "The dynamics of episodic recall"

### Supplementary Information

**a**

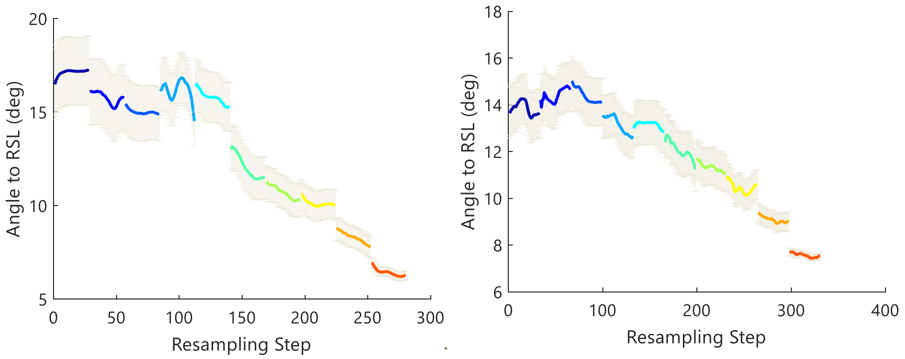


**Supplementary Fig. 1:** a. Angle to the RSL across all single-report events for the SAME (left) and DIFF (right) groups.
